# modelRxiv: A platform for the dissemination and interactive display of models

**DOI:** 10.1101/2022.02.16.480599

**Authors:** Keith D. Harris, Guy Hadari, Gili Greenbaum

**Affiliations:** Department of Ecology, Evolution and Behavior, The Hebrew University of Jerusalem, Jerusalem, Israel

## Abstract

Modeling the dynamics of biological processes is ubiquitous across the ecological and evolutionary disciplines. However, the increasing complexity of these models poses a significant challenge to the dissemination of model-derived results. With the existing requirements of scientific publishing, most often only a small subset of model results are generated, presented in static figures or tables, and made available to the scientific community. Further exploration of the parameter space of a model, investigation of possible variations of a model, and validation of the results in relation to model assumptions commonly rely on local deployment of code supplied by the authors. This can pose a technical challenge due to the diversity of frameworks and environments in which models are developed, and preclude model validation and exploration by readers and reviewers. To address this issue, we developed a platform that serves as an interactive repository of biological models, called `modelRxiv`. The platform provides a unified interface for the analysis of models that does not require any technical understanding of the model implementation. To facilitate adding models to `modelRxiv`, we utilize OpenAI large-language models (LLMs) to make code written in different programming languages compatible with `modelRxiv`, making the platform language-agnostic. `modelRxiv` is designed to serve as an interactive extension of published models, allowing users to regenerate model results under user-defined parameterizations of the model. By making published models accessible, this platform promises to significantly improve the accessibility, reproducibility, and validation of ecological and evolutionary models.

## 1 Introduction

Modeling the dynamics of ecological and evolutionary processes has been key to the development of ecology and evolution for more than a century [1]. Over the past few decades, the availability of computational power has paved the way for models of increasing complexity, involving numeric, stochastic, and individual-based modeling features, and exploration of multiple variables spanning over large parameter spaces [2]. Detailed and high-resolution investigation of models can offer important insights on general biological phenomena, as well as for specific systems. However, traditional scientific publishing standards—static papers depicting several model results representing select parameter values—are limited in demonstrating the full scope of model results and their implications. This affects both the dissemination of potential insights from a model and the ability to assess the robustness of model results during the scientific peer-review process.

Currently, models in ecology and evolution, and across the biological disciplines, are implemented in a number of different programming languages because there is no standard framework for publishing model implementations. Consequently, deploying models locally by a reviewer or a reader usually requires prior knowledge of the specific coding language or framework in which the model code was written, the installation of dependencies (e.g., code packages), and learning how to operate the model code and visualize results. This technical hurdle can make published models essentially inaccessible, where only the authors of the model are able to reproduce its results and alter the model.

Various platforms have been developed to address some of these difficulties. These approaches can be categorized into two broad classes: (i) a ‘model-centric’ approach, where the emphasis is on model analysis and visualization, and users are not required to interact with the underlying model code, and (ii) a ‘code-centric’ approach, where users interact with model code in order to analyze the model. Model-centric platforms provide a single programming language for modeling (e.g., *NetLogo* [3], *Mathematica* [4], *MATLAB* [5]), while code-centric platforms accommodate the deployment of multiple programming languages in a single interface (e.g., *Jupyter* [6], *CodeOcean* [7]). In both classes, models are accessed through a unified interface, reducing the investment needed by reviewers or readers to interact with the model. However, most published models are implemented in different programming languages, and translating model code so that it is compatible with a model-centric platform can be impractical. On the other hand, code-centric solutions that support multiple programming languages are focused on model implementation, and lack features that can make model analysis straightforward and accessible.

To bridge the gap between these two approaches, we developed an interactive repository of biological models called `modelRxiv` (https://modelrxiv.org). Our web-based platform allows users to visualize the results of many different types of published and unpublished models from the same interface without having to download or install framework-specific software. `modelRxiv` is designed with a simple user interface that does not require a technical understanding of programming or modeling to operate. To streamline the implementation of this protocol for existing models, and the translation of these models to browser-compatible programming languages, we provide an AI-assisted process of adapting the model code to the `modelRxiv` platform. We envision this platform as a tool for model exploration by reviewers and readers, and as a repository that can make published and unpublished models easily accessible to the scientific community.

## 2 Platform

`modelRxiv` is browser-based, with model computation and visualization occurring in the browser. The `modelRxiv` user interface has three main screens: (i) an index of models that can be filtered and sorted according to model metadata such as title, authors, model categories, and keywords (Fig. 1); (ii) model analysis pages, which include model visualization, regeneration of predefined model dynamics, and parameter manipulation features (Figs. 2 and 3); and (iii) a model upload page for uploading and updating models.

**Figure 1.**
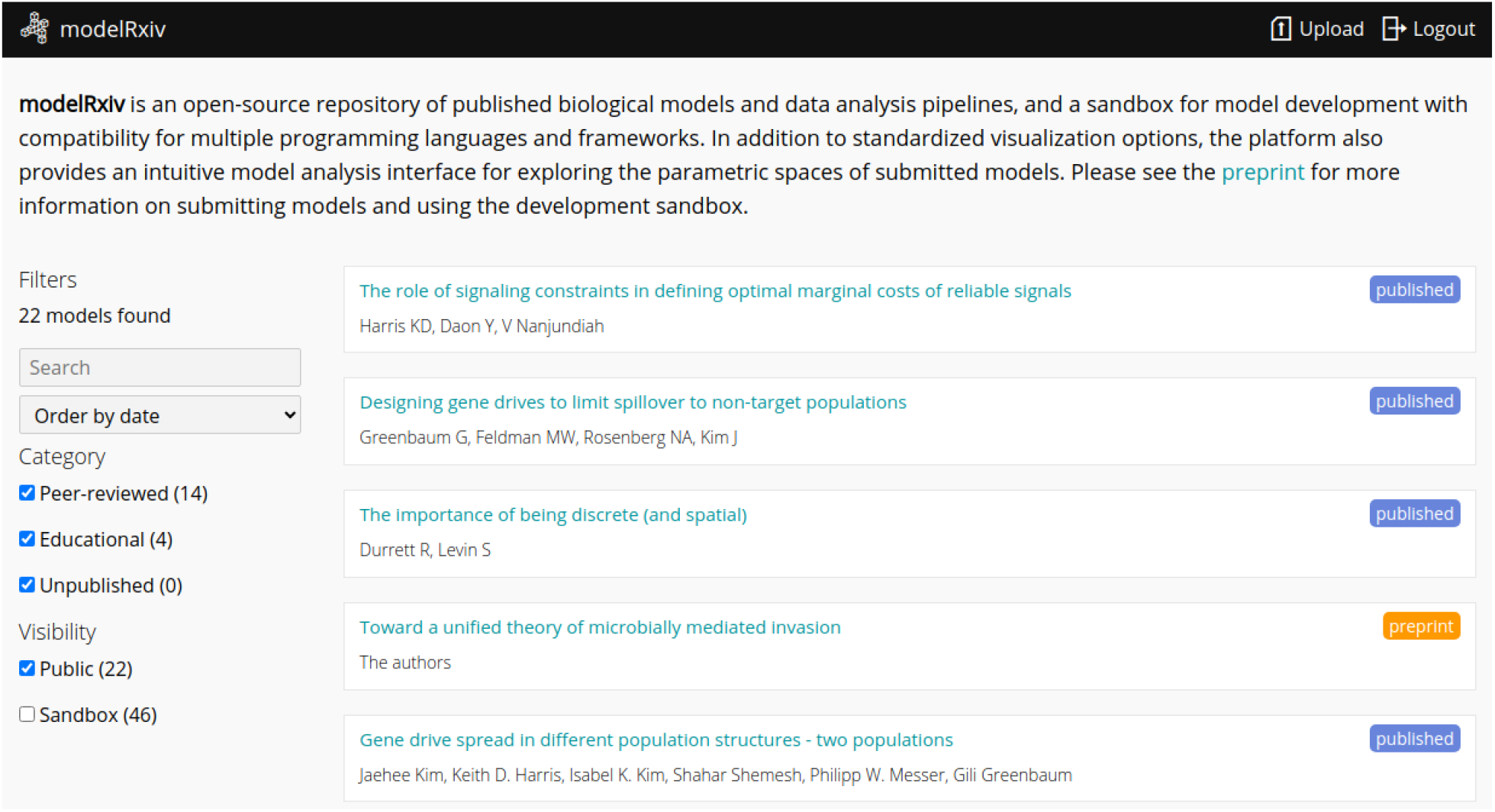
The main page of modelRxiv, which lists available models on modelRxiv belonging to different categories. On this page users can search and sort models, and select a model to view its ‘model analysis page’. When authenticated, the index will list models that are publicly available, as well as private models (‘Sandbox’ models) uploaded by the user.

**Figure 2.**
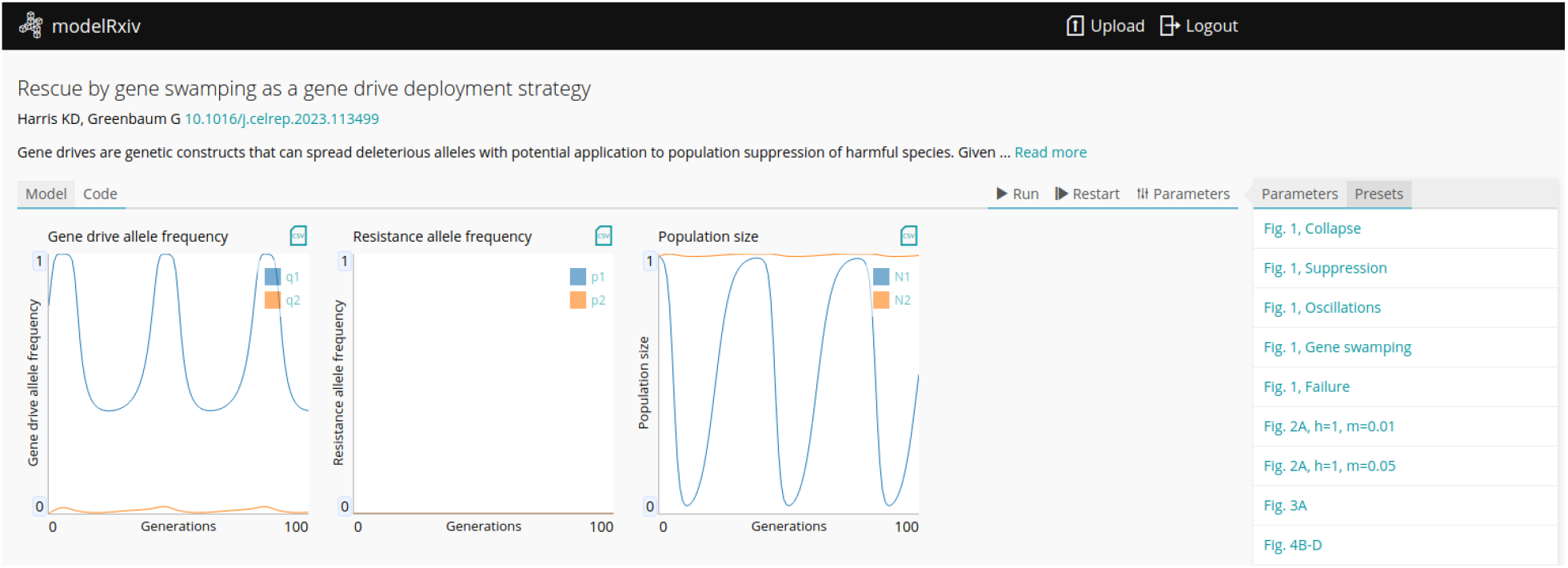
Dynamics of gene drive model generated on-the-fly. At the top of the page, metadata relating to the model is shown, including author information, the title and description of the model, and a link to the manuscript if available. Underneath are the dynamics that have been generated on-the-fly: in this case, the plots include the genetic and demographic dynamics of the gene drive model. On the right is the list of presets, which allows users to reproduce specific dynamics of the model, for example replicating the figures in the manuscript.

**Figure 3.**
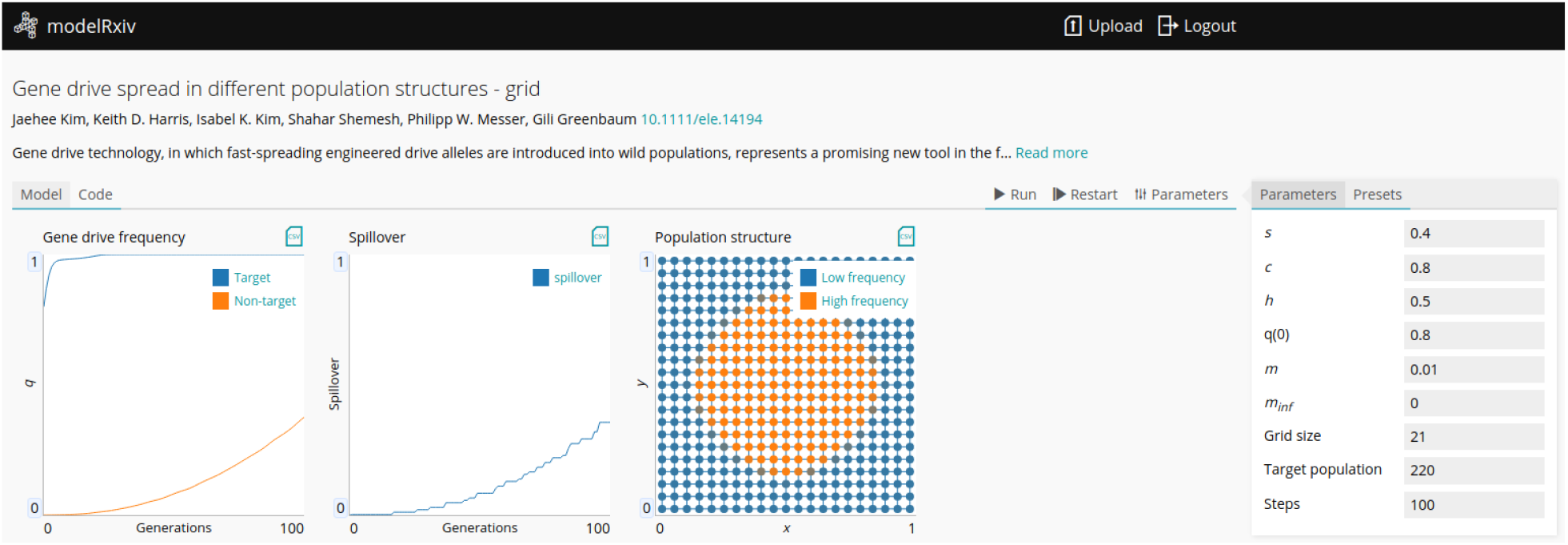
Spatial dynamics of a gene drive model that incorporates population structure and dispersal. The spread of the gene drive through populations is visualized as nodes with the color reflecting the frequency of the gene drive in each population (right panel). Users can track the overall spread as the number of populations above a certain threshold (the ‘Spillover’ panel) or observe the spatial dynamics (the ‘Population structure’ panel). These standardized displays can be used for many different types of models. Users can manipulate the parameters listed on the right and rerun the model dynamics to observe the effect of different parameter configurations on the result.

Models can be either ‘public’, viewable by everyone, or in ‘sandbox mode’ where only the owner can see and interact with the model. Public models can be viewed and analyzed without logging in or creating a user, but registration is necessary to upload models and analyze models in sandbox mode (registration is free and anonymous). In this section we describe the main features of `modelRxiv` that can be accessed through the model analysis page or the model upload page.

In order to explain the features of `modelRxiv`, we refer to published models that have been uploaded to `modelRxiv` that relate to gene drives (a genetic element that violates Mendelian inheritance) [8]. This field is an interesting use-case for `modelRxiv`, as having a repository of published gene drive models in a single platform could be instrumental in developing the interface between modelers and regulators [9–11].

### 2.1 Model analysis page

The model analysis page provides users with basic model metadata, such as publication information, as well as interactive figures that are generated on-the-fly by the user (Fig. 2). Users can interactively run model dynamics that will be visualized according to the type of output generated by the model (e.g., line plots for continuous variables, 2-dimensional grid for visualizing spatial dynamics).

Model exploration can be conducted either by regenerating results of parameter sets specified by the model authors, or by specifying custom parameters. Models uploaded to `modelRxiv` can have multiple parameter ‘presets’, meaning parameters that relate to a certain scenario of interest in the model (Fig. 2). These can be linked to a figure in a published manuscript, allowing users to replicate published figures, and continue exploring the model by examining the effect of changing model parameters while starting off the exploration from the published parameters. Presets make model results more accessible to users, by guiding them to specific parameter sets that generate dynamics of interest. The presets menu is opened by default on the right side of the page if presets are provided by the authors. Clicking on preset labels will run model dynamics for the specified preset. Users can also control the dynamics with the control buttons at the top right corner of the model analysis page. Models can be run by clicking the ‘Run’ button, and can be stopped by clicking ‘Pause’. Clicking ‘Restart’ will end the current model dynamics and restart with the currently selected parameters (Fig. 2). If users wish to generate additional visualizations of the dynamics, or aggregate outputs, results generated by the model analysis page can be exported as CSV files by clicking the ‘CSV’ icon at the top right corner of the plot.

In addition to running parameter presets, users can manipulate models parameters and rerun analyses, testing the effect of different parameter values on the results (Fig. 3). Clicking the ‘Parameters’ tab on the right side of the model analysis page will open a list of parameters that are available and can be modified; after changing parameter values, users can rerun the model dynamics by clicking ‘Run’ or ‘Restart’ (Fig. 3).

Model authors and users can also run meta-analyses of the model, where the model is run repeatedly with different parameter sets to produce a more complex summary plot. These meta-analyses can be defined using the model scheme, to indicate which function should be run in order to generate the data, and what plot should be used; model computation will be automatically distributed in the browser. For instance, for the gene drive model, we can generate a grid of results by running the model for a grid of *s* and *c* values, and defining a color for each result. This result is then plotted in the browser using a predefined plot type for grids.

### 2.2 Uploading models

To allow flexibility in the integration of models with `modelRxiv`, we developed a language-agnostic protocol for model inputs and outputs. This protocol requires the implementation of a single step function in order to integrate with `modelRxiv`, in addition to a text file describing the model and its parameters. We also developed an AI-assisted pipeline to convert models to be compatible with this protocol.

To streamline the process of uploading models to `modelRxiv`, we developed an upload interface that allows users to upload their code and validate that it is compatible with `modelRxiv`. The upload form is accessible to authenticated users by clicking the ‘Upload’ button at the top right corner of the screen. The process is separated into two main text fields: (i) a scheme describing the model metadata, parameters, plots and presets; and (ii) a text box containing the model code (the step function and any other functions required by the model).

The upload form also includes AI-assisted features to help users make their code compatible with `modelRxiv`. These features utilize a number of OpenAI large-language models LLMs [12]. This feature is made feasible by the simple design of our model protocol, as well as the model scheme pseudo-code, which is designed to balance readability by users and LLMs, and be easily processed by `modelRxiv`. Instructions and examples are provided to the LLM alongside user requests to ensure that generated code is compatible with `modelRxiv`. Additionally, specific OpenAI models can be fine-tuned with training and validation datasets in order to improve the accuracy of responses to prompts. For the current version, we used models already added to `modelRxiv` to fine-tune the LLMs used for scheme generation and model conversion.

As models are added to `modelRxiv`, we can further improve these pipelines, adding new publicly available models to the training dataset.

To utilize these features, users paste their model code into the code editor, and click the ‘Generate’ button (Fig. 4). This will first generate a model scheme if one has not been created, based on the parameters that can be manipulated in the code. After generating a scheme, users can convert the code to be compatible with `modelRxiv` by clicking the ‘Convert’ button. This will use another LLM to convert the code to a language compatible with `modelRxiv` if applicable, and ensure it implements the ‘step’ function correctly, and that parameters correspond to those present in the model scheme. The result can then be validated within the form using the ‘Test’ button. If the test fails, users can send the code along with the error to the AI assistant; this will generate a new version of the code or scheme addressing the error. In addition to technical validation of compatibility, model dynamics should be manually validated against the original outputs to assure that the model has been imported correctly. In our development of this pipeline, we successfully converted model code from a variety of commonly-used modeling languages, including *MATLAB, Mathematica*, and *R*.

**Figure 4.**
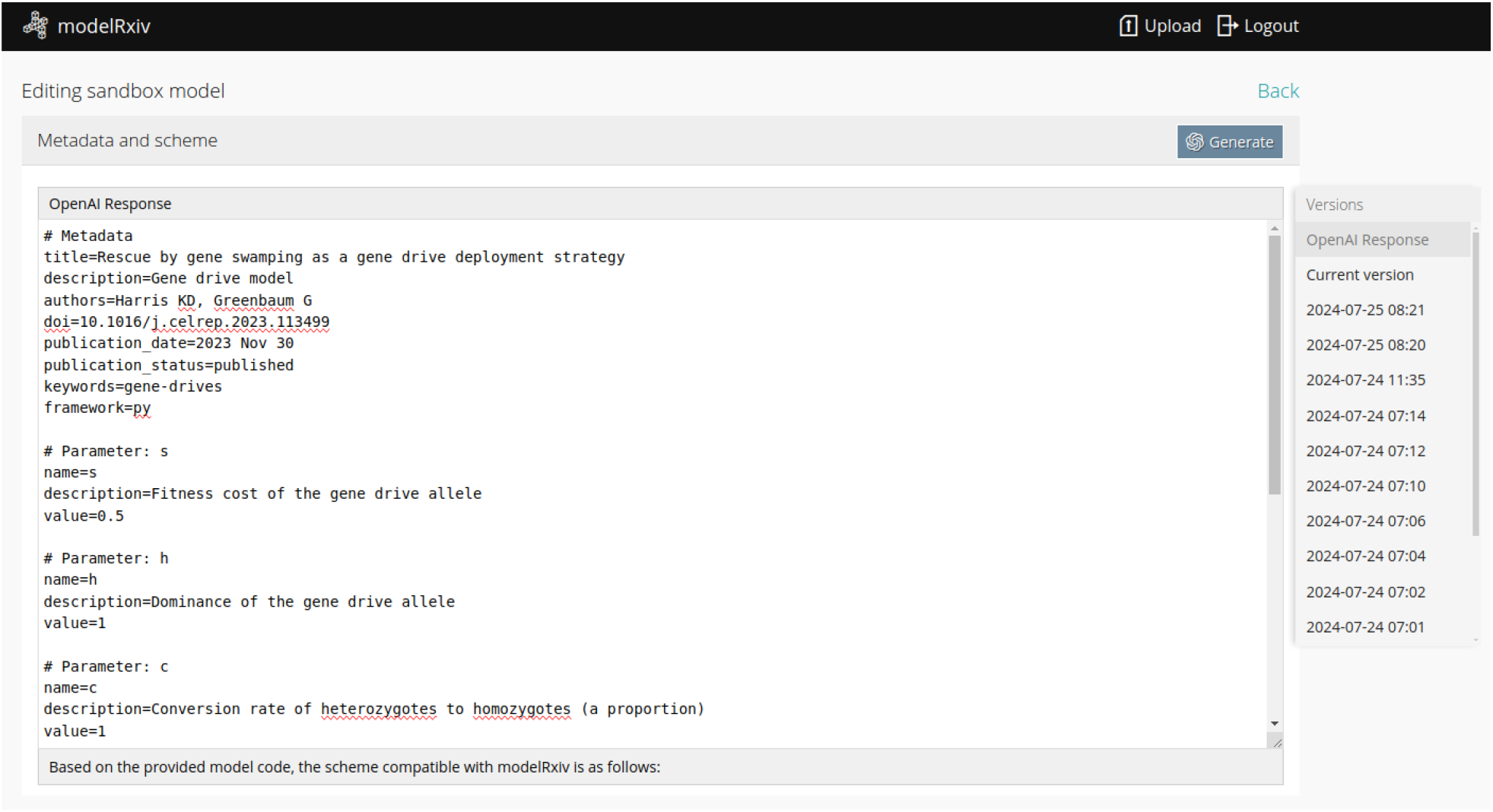
Upload interface with scheme code generated by LLM. The first section of the model editing form is the model scheme. In this example, a model scheme was generated with the AI-assistance feature, by providing basic metadata of the model and the model code. The response of the LLM appears as a new tab in the editor; previous versions of the scheme or code can be loaded by clicking on the version in the ‘Versions’ menu on the right.

After uploading by clicking the ‘Submit’ button at the end of the upload form, models are visible and accessible only to the account owner, and appear as ‘Sandbox’ models. This gives users the ability to ensure that the model operates correctly before submitting it to the public `modelRxiv` repository. Uploaded models can be edited using the ‘Edit’ button on the model page. Model code and parameters are versioned, so that previous versions can be browsed and restored if necessary. To submit the model to the public repository, users click the ‘Publish’ button on the model analysis page. This will transfer the model to a moderator who will review the model manually to ensure it is valid, and subsequently the model will be publicly available on `modelRxiv`.

## 3 Discussion

Here we presented and described the features of the `modelRxiv` platform, an interactive repository of models. The purpose of the platform is to facilitate reviewers and readers in evaluating and investigating models, to increase the accessibility of published and unpublished models in ecology and evolution, and to serve as an educational tool. To allow a common interface for multiple types of models, we developed a language-agnostic protocol for adding models to `modelRxiv`, and integrated a number of customized LLMs into the process of adding models to the repository. These LLMs are able to adapt models written in various programming languages to languages supported by `modelRxiv`, and ensure compatibility with the `modelRxiv` protocol. This interface includes validation of model code for compatibility errors. These features remove many of the technical hurdles that can be encountered in adapting code to specific frameworks of model-centric platforms.

In our development of the model uploading pipeline, we found current LLMs capable of converting a variety of programming languages to `modelRxiv` without introducing errors into the code. In most cases, the LLM managed to convert model code to Python code compatible with `modelRxiv` in a single step. The examples we used were often complex ecological models that included many parameters. The successful integration of generative AI features into `modelRxiv` relies on a number of features of our platform: (i) the model protocol is straightforward and identical for all model types; (ii) the model scheme we developed is robust to typographical errors and can be read and generated by LLMs; (iii) using Python as a default language makes model conversion simpler, since LLMs contain extensive Python-related information in their training data; and (iv) models can be validated easily owing to the use of unified outputs. LLMs are particularly suited to cases where the output pattern has well-defined guidelines that can be enhanced by providing previous examples. Thus, `modelRxiv` is a perfect use-case for LLM-assisted coding. In addition, our platform will be able to use the growing repository of models in `modelRxiv` to fine-tune the AI-assisted conversion of models; we have already implemented fine-tuning for our current version based on the models already added to the platform.

With the development of new platforms such as `modelRxiv`, it is important to evaluate and compare the main features to existing platforms to better understand their potential contribution. There are a number of established comparable platforms that provide an interface for manipulating model parameters and visualizing dynamics or results, mainly *Mathematica, MATLAB, Jupyter* and *NetLogo*. When designing `modelRxiv`, we considered the features already offered by these platforms, summarized in Table 1.

**Table 1:** Features of modelRxiv compared to other platforms that are used for modeling. We define three broad categories to differentiate between platforms: (i) the “approach” of the platform, whether it is primarily designed for models or for code; (ii) the modeling functionality offered, whether the platform allows model analysis or is a repository; and (iii) where computation occurs, which determines what costs and technical hurdles are involved in analyzing models. For the computation category, “client” indicates that the end-user is responsible for providing resources for computation, either by running a program locally (not in the browser) or by using a paid cloud service.

| Platform | Approach | Functionality | Computation in browser? | Language-agnostic? |
| --- | --- | --- | --- | --- |
| **modelRxiv** | Model-centric | Repository/Analysis | Yes | Yes<sup>d</sup> |
| *NetLogo* [3] | Model-centric | Repository/Analysis | NetLogo Web<sup>a</sup> | No |
| *Mathematica* [4] | Model-centric | Analysis | No | No |
| *MATLAB* [5] | Model-centric | Analysis | No | No |
| *Numerus* [13] | Model-centric | Analysis | Export to Web<sup>b</sup> | No |
| *EBI BioModels* [14] | Model-centric | Repository | Not applicable<sup>c</sup> | Yes |
| *Jupyter* [6] | Code-centric | Analysis | No | Yes |
| *CodeOcean* [7] | Code-centric | Code validation | No | Yes |

<sup>a</sup> *NetLogo* also has a browser-based version that includes a library of models suitable for running in a browser.
<sup>b</sup> *Numerus* models can be exported to a format that can be run in a browser.
<sup>c</sup> *BioModels* contains previously generated plots and is not designed for model analysis.
<sup>d</sup> By providing an AI-assisted pipeline for adding models, **modelRxiv** can accept as input models written in many different frameworks.

To make a repository of models accessible to a broad audience, it is crucial that models can be interacted with easily, without requiring the user to install additional tools or understand the specific programming language used. Indeed, some modeling platforms offer web-based implementations as an additional feature, where model analysis and visualization occurs in the browser. In this type of web-based setup, the user is not required to deploy computational containers, making the service sustainable (Table 1). However, these features are not widely used, and are often not feasible because most modeling languages lack browser support. The adoption of a repository depends on the majority of models in a field being compatible with it, but integration of existing models with platforms that are based on specialized programming languages is a substantial challenge. On the other hand, ‘code-centric’ platforms that support execution of multiple programming languages, such as *Jupyter* [6] or Code Ocean [7], often require deployment to cloud resources. The code-centric design of these platforms can also introduce technical hurdles unrelated to understanding the models themselves. Thus, a platform such as `modelRxiv` that accepts as input models written in different languages, while providing an interface that is designed specifically for modeling, could bridge the gap between model-centric and code-centric approaches, making accessible many different types of models in a single repository.

### 3.1 Facilitating evaluation of models during and after the review process

One of the most important potential applications of `modelRxiv` is to facilitate and improve the review processes of modeling studies in ecology and evolution. During the review of modeling studies reviewers should, ideally, evaluate the correctness, robustness, reproducibility, and applicability of models. However, in practice, even when the model code is attached to the submission, there are technical difficulties including setup, installation of dependencies, compatibility, and acquaintance with the specific coding language chosen by the author that makes this time-consuming and unfeasible in almost all cases. Consequently, models are rarely reproduced by reviewers during the review process, and are even more rarely subjected to manipulation and thorough investigation beyond the specific parameter values chosen by the author.

Improving model evaluation during the review process requires development of user-friendly and accessible tools for reviewers. The understanding that such tools are vital to ensure code validation has led to the adoption of services for deploying data processing code to computational containers by reviewers, such as *CodeOcean* [7, 15]. *CodeOcean* provides an interface through which reviewers can manipulate and deploy code associated with a manuscript during the review process. By removing the technical hurdles of code deployment, code validation can become an integral part of the review process without resulting in a significant additional burden on reviewers.

`modelRxiv` can facilitate testing model robustness beyond simply validating model code for technical errors. With models uploaded to `modelRxiv`, reviewers can extensively explore the full parameter space of the model, and can even alter the model itself. Using the presets feature, reviewers could easily generate figures from the manuscript, and assess the robustness of the model in terms of the parameters used to generate these figures. By using presets, the model owner can guide users through stages of exploring the parametric space of the model. This can lead to a more intuitive understanding of the model than a static figure, as the user is free to manipulate the model parameters and observe the effect on the model result, or to visualize model dynamics for different regions within a certain figure.

Making model code validation an integral part of the review process would improve not only confidence in the model results, but also encourage more thorough exploration of the model parameter space and the underlying assumptions by the authors. It could also shorten the review process by allowing reviewers to answer queries regarding model parameters without having to rely on the authors to produce additional figures.

As `modelRxiv` is a public repository, using it in the review process has the important benefit of ensuring that the model would be accessible to readers of the manuscript after it has been published. This may encourage readers of the paper to “play” with the model, potentially leading to new insights and directions of research.

### 3.2 Educational uses

`modelRxiv` also serves as an educational tool. With models that demonstrate basic principles in ecology and evolution, students can visualize pre-prepared dynamics and manipulate model parameters to gain intuition on the phenomenon in question, and they can test hypotheses regarding the relationship between model parameters. At a more advanced level, students can explore the code implementation of the model, to generate similar alternative models, and to gain experience in model coding and design. The fact that the user is not exposed to the full complexity of the model from the very beginning is an important aspect when encouraging those not familiar with model design or programming to engage in exploration of the model. Therefore, the clear separation of model visualization and manipulation from the underlying code would be helpful in teaching environments, where it is necessary to account for variable technical abilities of students to provide individualized and flatter learning curves for students.

### 3.3 Public accessibility of model results

Beyond facilitating the review process, we believe that `modelRxiv` could promote and improve the exploration of published models in the ecological and evolutionary disciplines. First, the ease of accessibility to published models would incentivize researchers to expand existing models and utilize existing modeling frameworks, thereby making model development faster. This would also improve the comparability between published models, ensuring that conclusions pertaining to differences between model results can be coherently attributed to changes in key assumptions, rather than to model design or coding. Second, encouraging the scientific community to participate in manipulation of model parameters, in a deeper examination of model parameter spaces, and in adjustment of model assumptions through simple alteration of underlying code, could generate new insights for existing models. Such inquiries and modifications could lead to the identification of novel model behaviors with biological significance, perhaps undetected due to a different focus of the original study. These investigations could also lead to a deeper understanding of the model behaviors, particularly in terms of the boundaries in which the described behaviors of the models no longer hold, and a discussion on the biological significance of these boundaries. Therefore, adoption of `modelRxiv` by the eco-evolutionary modeling community could encourage collaborations between researchers to extend and elaborate on modeling studies, for example between the publishers of the original modeling study and the researchers identifying interesting behaviors in their models.

## 4 Methods

### 4.1 Implementation of platform

`modelRxiv` is browser-based, and was designed as a serverless application. This greatly reduces operational costs as there is no need to operate a back-end server, and costs scale with use. These considerations are important to make the project sustainable in the long-term as a free, open-source repository. Further technical details of the implementation are available on the GitHub repository of the project (https://github.com/carrowkeel/modelrxiv).

### 4.2 Model protocol

To allow `modelRxiv` to act as a wrapper for models, we developed a protocol with minimal requirements for model integration. This protocol is specifically designed for step-wise models, but will be suitable for any model that generates results through iteration. Model code must implement a ‘step’ function that contains the main model logic, in addition to any other functions that were part of the model code.

In addition to the model code, we designed a model scheme pseudo-code that balances readability by users editing the scheme manually, generation and modification by LLMs, and processing by `modelRxiv`. The scheme is separated into sections with each section describing a different aspect of the model, including metadata, parameters, plots and presets. The AI-assisted model uploader can generate model schemes based on code, and attempt to complete or correct partially written schemes.

At present, `modelRxiv` directly supports code written in Python or JavaScript, as these can be easily executed in a browser environment. However, models written in other languages can be integrated with `modelRxiv` by using the AI-assisted conversion pipeline; the simplicity of the model protocol makes this pipeline feasible.

## 5 Accessibility of data

The `modelRxiv` platform and all models presented here are freely accessible at https://modelrxiv.org. The platform code is available at github.com/carrowkeel/modelrxiv and github.com/carrowkeel/apc. All platform code is licensed under AGPLv3. Models are licensed under CC-BY 4.0 unless otherwise stated. The OpenAI models used in this version of `modelRxiv` are `gpt-4o` and fine-tuned versions of `gpt-4o-mini`. To allow users to provide feedback on bugs and issues using `modelRxiv`, we opened a *Slack* workspace that can be joined using the invitation link on the ‘contribute’ page (https://modelrxiv.org/contribute.html).

## 6 Author contributions

KDH designed and developed the platform, GG supervised the development of modeling features, and GH consulted on cloud integration. KDH and GG wrote the manuscript. All authors read and approved the final version of the manuscript.

## 7 Acknowledgments

We would like to thank Shachar Perlman and Jonathan Halperin from the Alpha Program in The Hebrew University Youth Division for testing features of the platform, and David Gokhman, Oren Kolodny and members of the Greenbaum Lab for helpful comments and discussions. This project was supported by Israel Science Foundation (ISF) Grant 2049/21, and by German-Israel Foundation (GIF) Grant I-1526-500.15/2021.

## References

1. Otto, S. P. & Day, T. A biologist’s guide to mathematical modeling in ecology and evolution. Princeton University Press (2007).

2. Grimm, V. & Berger, U. Structural realism, emergence, and predictions in next-generation ecological modelling: Synthesis from a special issue. Ecological Modelling 326, 177–187 (2016).

3. Tisue, S. & Wilensky, U. Netlogo: A simple environment for modeling complexity. International Conference on Complex Systems 21, 16–21 (2004).

4. Wolfram, S. Mathematica: a system for doing mathematics by computer. Addison Wesley Longman Publishing Co., Inc. (1991).

5. MATLAB. MATLAB 9.11 (R2021b). The MathWorks Inc. (2021).

6. Kluyver, T. et al. Jupyter Notebooks-a publishing format for reproducible computational workflows. IOS Press (2016).

7. Staubitz, T., Klement, H., Teusner, R., Renz, J. & Meinel, C. CodeOcean-A versatile platform for practical programming excercises in online environments. 2016 IEEE Global Engineering Education Conference, 314–323 (2016).

8. Bier, E. Gene drives gaining speed. Nature Reviews Genetics 23, 5–22 (2022).

9. Long, K. C. et al. Core commitments for field trials of gene drive organisms. Science 370, 1417–1419 (2020).

10. Combs, M. A. et al. Leveraging eco-evolutionary models for gene drive risk assessment. Trends in Genetics 39, 609–623 (2023).

11. Frieß, J. L., Lalyer, C. R., Giese, B., Simon, S. & Otto, M. Review of gene drive modelling and implications for risk assessment of gene drive organisms. Ecological Modelling 478, 110285 (2023).

12. OpenAI. Introducing GPT-4 https://openai.com/blog/gpt-4. Accessed: 2024-01-01. 2023.

13. Getz, W. M., Salter, R., Muellerklein, O., Yoon, H. S. & Tallam, K. Modeling epidemics: A primer and Numerus Model Builder implementation. Epidemics 25, 9–19 (2018).

14. Le Novere, N. et al. BioModels Database: a free, centralized database of curated, published, quantitative kinetic models of biochemical and cellular systems. Nucleic Acids Research 34, D689–D691 (2006).

15. Cheifet, B. Promoting reproducibility with Code Ocean. Genome Biology 22, 65 (2021).

